# Graded NTCP dynamics: leveraging patient-reported outcomes to explore response and cross-toxicity trade-offs in head and neck cancer

**DOI:** 10.1101/2025.10.13.682060

**Authors:** Daniel James Glazar, Ryan Werthmann, Niema B. Razavian, Renee Brady-Nicholls

## Abstract

1

**Background:** Radiotherapy (RT) is a localized therapy used to treat approximately 50% of all cancer patients and 75% of head and neck cancer (HNC) patients. Despite tumor response to RT, off-target effects to surrounding organs at risk (OAR) can result in toxicities that exacerbate patient symptoms and necessitate interventions such as tube feeding. While current models of normal tissue complication probability (NTCP) can account for complex OARs-symptoms interactions, they only allow for dichotomous outcomes without consideration of graded response. Polytomous patient-reported outcomes (PROs), collected via patient surveys, capture patients’ experience of their symptoms throughout treatment. Unfortunately, such measures are rarely incorporated when modeling treatment-related toxicities.

**Purpose:** In this study, we aimed to develop a model of NTCP to recapitulate population-level PRO dynamics from a published dataset and explore response and cross-toxicity trade-offs in HNC.

**Methods:** We employed the classical linear-quadratic dose-response model to describe tumor response to RT with logistic growth. As a surrogate for normal tissue damage, we modeled absorbed dose kinetics to individual OARs (buccal mucosa, oral cavity, superior pharyngeal constrictor muscle (sPCM), and body) using a one-compartment pharmacokinetic model with linear elimination. We then employed a minimal inhomogeneous continuous-time Markov chain model (with tumor size and absorbed dose as time-varying covariates) to describe PRO dynamics (oral pain, dysphagia, weight loss, and tube feeding, measured on the EORTC-HN35 scale). We modeled symptom-specific NTCP as the cumulative incidence of severe symptom. Finally, we explored various response-toxicity and toxicity-toxicity trade-offs with respect to dose, OAR sparing, and fractionation.

**Results:** The developed mathematical model recapitulated the qualitative features of the motivating published dataset, including 1) a transient reduction of a subset of symptoms for 1–2 weeks; followed by 2) an acute exacerbation of symptoms throughout the rest of RT; followed by 3) a long-term relaxation of symptoms to below baseline levels.

Response-toxicity trade-offs were sensitive to dose, variably sensitive to OAR sparing (dependent on OAR-symptom associations), and sensitive to fractionation. Toxicity-toxicity trade-offs were insensitive to dose, sensitive to OAR sparing, and insensitive to fractionation.

**Conclusions:** Overall, this mathematical model addresses some of the limitations of current NTCP models by explaining graded toxicity dynamics. By integrating both tumor control and quality of life considerations into a singular model, recommendations of dose, OAR sparing, and fractionation can be made. Future iterations of the model could aid clinicians in RT dose-finding and selecting a RT plan that will optimize tumor control and patients’ quality of life.

## 2 Introduction

Radiotherapy (RT) is an effective oncologic treatment that is used to treat approximately 50% of all cancer patients^1^ and 75% of head and neck cancer (HNC) patients.^2^ Despite its utility, off-target effects are common within and adjacent to the RT field. The resulting damage to normal healthy tissues exacerbates patient symptoms and causes treatment-related adverse events (trAE) that impact patients’ quality of life. For example, in HNCs irradiation of the buccal mucosa, oral cavity, and superior pharyngeal constrictor muscle (sPCM) is associated with mucositis, oral pain, and dysphagia, which can result in significant weight loss that requires feeding tube insertion.^3,4^ While limiting RT dose to the sPCM may help avoid dysphagia, it comes with a trade-off: either at the expense of tumor coverage or at the expense of normal tissue toxicity (e.g., oral pain).

To measure trAEs, the US National Cancer Institute (NCI) has developed the Common Terminology Criteria for Adverse Events (CTCAE) for clinicians. Unfortunately, these reports often lack reliability^5^ and underestimate patient symptoms.^6–8^ Consequently, the patient-reported outcomes (PRO) version of the CTCAE (PRO-CTCAE) was developed to more directly assess patient symptoms.^9,10^ Other PRO instruments have been subsequently developed to measure general (e.g., FACT-G,^11^ EORTC-QLQ-C30^12^) as well as disease-specific (e.g., FACT-HN^13^, EORTC-QLQ-HN43^14^) and treatment-specific (e.g., FACT-HN-RAD^15^) symptoms.

One clinical question of interest remains: how should trade-offs between tumor response and multiple treatment toxicities be balanced? To address this question, ^161718^mathematical models of tumor control probability (TCP)^19,20^ and normal tissue complication probability (NTCP)^4,21,22^ models have been developed. Perhaps the most popular NTCP model is the Lyman-Kutcher-Burman (LKB) model. However, the LKB model assumes that only a single OAR is implicated in the symptoms of interest. This doesn’t account for the complex relationships between organs at risk (OARs) and symptoms. To address this, logistic regression models allow for more elaborate relationships between symptoms and OARs. Other, more elaborate models have subsequently and continue to be developed to account for the microscopic biological mechanisms underlying tumor response dynamics and normal tissue damage (e.g. damage and repair processes,^16^ oxygen depletion and reoxygenation,^17^ and cancer stem cell radioresistance).^18^ However, to our knowledge none of these models can explain graded symptom dynamics.

To this end, a variety of approaches have been and continue to be explored. For example, a dynamic factor model joined with a Ornstein-Uhlenbeck process has been employed to explain PRO dynamics using latent symptom constructs.^23^ In another study, an inhomogeneous continuous-time Markov chain (ICTMC) model joined with a tumor growth inhibition (TGI) model has been employed to describe the joint dynamics of PROs and tumor response.^24^ In a third study, PRO dynamics have been correlated to tumor response dynamics and clinical outcomes by looking at their cumulative change relative to baseline.^25^

Herein, we present a mathematical model to describe PRO dynamics and explore response-toxicity as well as toxicity-toxicity trade-offs in HNC patients undergoing RT.^26^ We model tumor response using the classical linear-quadratic (LQ) model with logistic growth.^27,28^ For toxicity, we model absorbed dose kinetics as a surrogate for normal tissue damage using a one-compartment pharmacokinetic model with linear elimination. We then employ an ICTMC model with tumor size and absorbed dose as time-varying covariates. NTCP is then modeled as the cumulative incidence of severe symptom. Finally, we explore the effects of dose, OAR sparing, and fractionation on response-toxicity and toxicity-toxicity trade-offs.

## 3 Materials and methods

### 3.1 Motivating data

Data motivating this work come from a prospective study of 859 HNC patients treated with RT between 2007 and 2017 at the University Medical Center Groningen in The Netherlands.^29^ In this study, patients with early larynx, infrahyoideal, or suprahyoideal cancers received 70 Gy in 2 Gy fractions (with or without concurrent systemic therapy) delivered through modern radiotherapy techniques (IMRT and/or VMAT). Patients completed the European Organization for Research and Treatment of Cancer (EORTC) Quality of Life Questionnaire–Head and Neck Cancer module (QLQ-HN35) at regular intervals during and up to 5 years after treatment.

For this study, we focused our attention on four treatment-related toxicities that were elevated in the patients with suprahyoideal (oropharynx, oral cavity, nasopharynx) HNCs: oral pain, dysphagia, weight loss, and tube feeding. All four symptoms were measured on the EORTC QLQ-HN35 scale. Each of these toxicities are associated with RT dose delivered to particular organs at risk (OAR; Figure 1A–B):^3,4^ oral pain (often due to mucositis) with the buccal mucosa and oral cavity, dysphagia with the oral cavity and sPCM, and weight loss with the sPCM and whole body integral dose (defined as the mean dose delivered to the patient normalized to the volume of the patient’s body). Additionally, tumor size has been implicated in oral pain and weight loss. Finally, insertion of a feeding tube and treatment breaks are necessitated when the burden of the preceding three symptoms becomes too high. For simplicity, in the modeling we assume that symptoms are independent conditioned on tumor size and absorbed dose kinetics to the associated OARs (Figure 1B). We also model tube feeding as conditioned on tumor size and absorbed dose kinetics rather than directly on other symptoms for simplicity.

**Figure 1.**
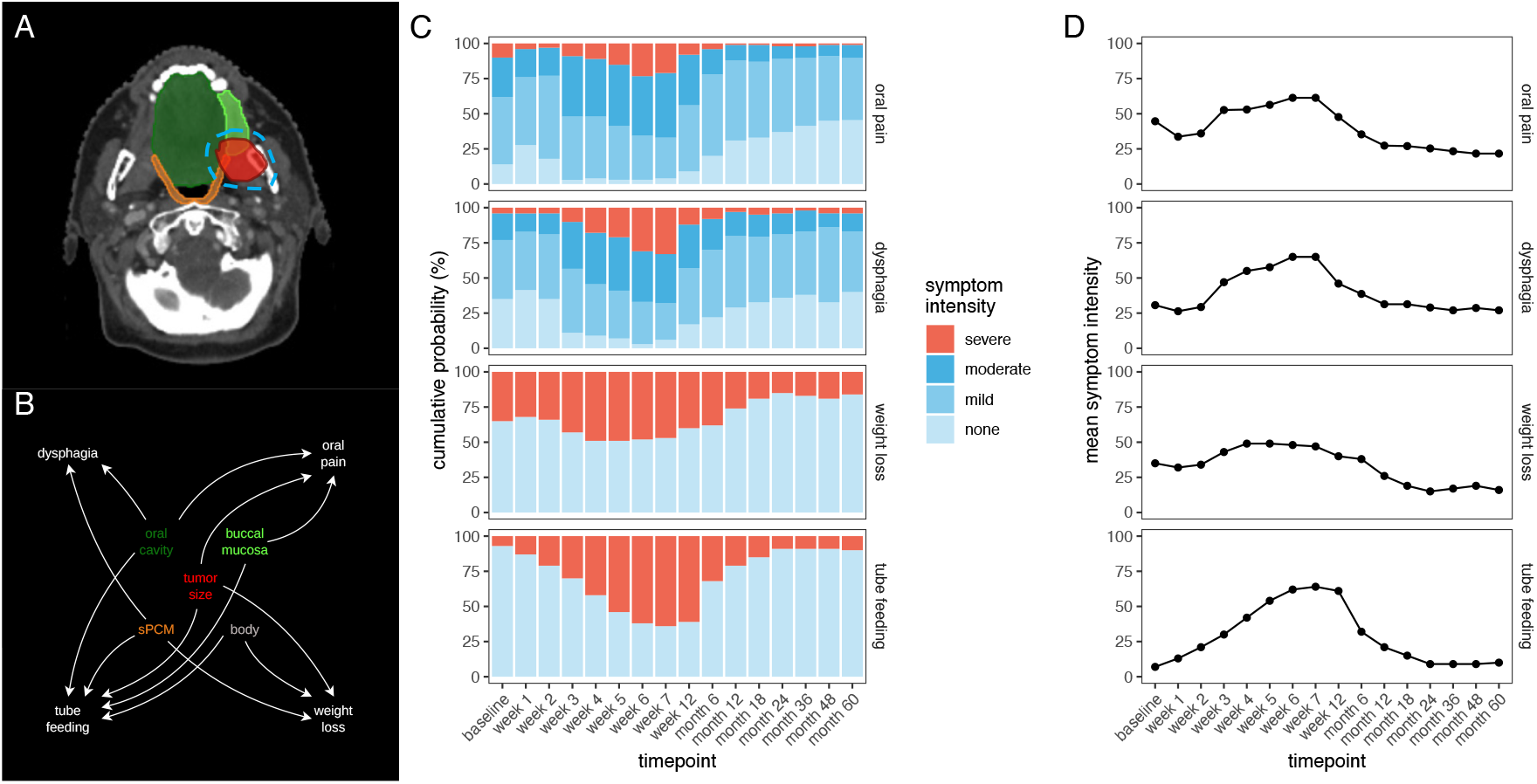
Motivating data. **A)** Anatomical locations of GTV and OARs implicated in symptoms from Brouwer, *et al*.^39^ Screenshot taken from *eContour* (http://econtour.org/cases/72; image 164/339).^40^ The original dataset is disease-free, and GTV is drawn here for demonstration purposes. PTV isodose line (blue) is drawn to maximize tumor response without consideration of OARs. Anatomical labels are in **B)** OAR-symptom associations. **C–D)** Population-level PRO dynamics over 60 months from Van den Bosch, *et al*.^29^ **C)** Symptom intensity probability distribution. **D)** Mean symptom intensity, ranging from 0 (none) to 100 (severe).

Oral pain and dysphagia scores were discretized onto a 4-point Likert scale (none, mild, moderate, severe), whereas weight loss and tube feeding had binary response (i.e., presence or absence). The population-level PRO dynamics for these symptoms are shown in Figure 1C. The scaled mean symptom severity, ranging from 0 (none) to 100 (severe), for each symptom are shown in Figure 1D.

### 3.2 Modeling tumor response to radiotherapy

Following Zahid et al.,^30^ we model tumor response to RT using the classical linear-quadratic (LQ) dose-response model^27^ with logistic growth (Figure 2A):

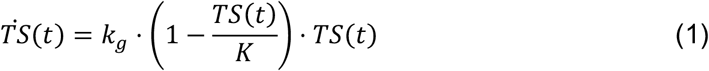

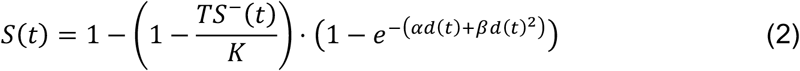

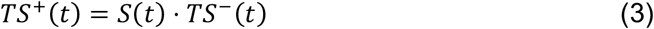

where *TS*(*t*) is the tumor size at time *t* [day]; *k_g_* is the tumor growth rate [day^-1^]; *K* is the tumor carrying capacity; *S*(*t*) is the tumor survival fraction upon administration of radiation dose at time *t*; *α, β* are radiosensitivity parameters [Gy^-1^, Gy^-2^]; *d*(*t*) is the radiation dose [Gy] delivered to the tumor at time *t*; and *TS*^−^(*t*), *TS*^+^(*t*) are tumor size immediately before and after administration of radiation at time *t*, respectively.

**Figure 2.**
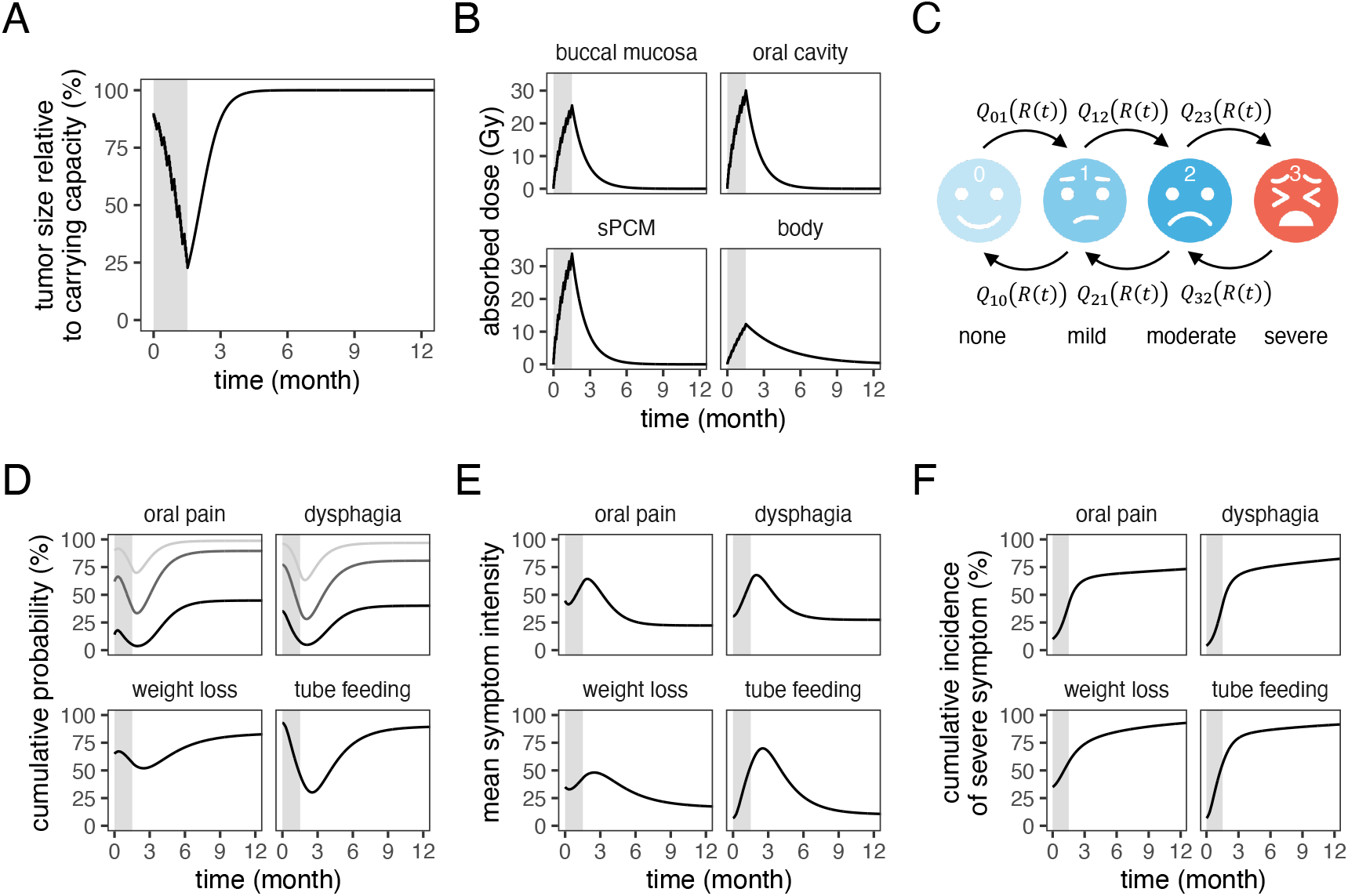
Simulated tumor response and toxicity dynamics. A simulated patient was treated with 70 Gy to the tumor with maximum tumor control and using standard fractionation (2 Gy per fraction for 35 fractions over 7 weeks (gray region)). **A)** Tumor response dynamics. **B)** Absorbed dose kinetics for each OAR. **C)** Schematic of Markov chain model for PROs. **D)** Symptom intensity probability distribution. Black, dark gray, and light gray curves refer to probability of no, mild, and moderate symptom for oral pain and dysphagia. Black curve refers to no symptom for weight loss and tube feeding. **E)** Mean symptom intensity, ranging from 0 (none) to 100 (severe). **F)** Cumulative incidence probability of severe symptom. mathematical models of tumor control probability

### 3.3 Modeling treatment-related adverse events

#### 3.3.1 Absorbed dose kinetics

Although RT uses high-energy x-rays to affect an acute tumor response, off-target biological effects accumulate over the course of treatment to induce trAEs.

Therefore, as a surrogate for long-term cumulative damage to and repair of normal tissue, we modeled absorbed dose kinetics *A_i_*(*t*) [Gy] to anatomical structure *i* (PTV, OAR, or other peripheral tissue) using a one-compartment pharmacokinetic model with bolus dose *d_i_* (*t*) delivered to anatomical structure *i* and a constant first-order elimination rate *k_El_,_i_* [day^-1^] (Figure 2B). For readability, we omit the subscript *i*.

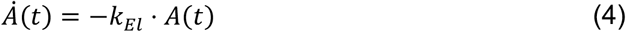

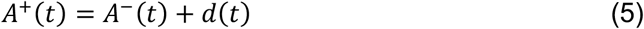

Solving equations 4 and 5 explicitly then gives us

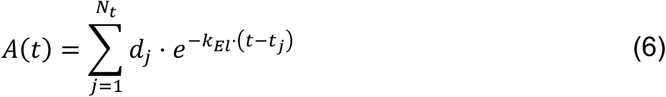

for RT dose *d_i_*, administration *j* at time *t_i_*, with *N_t_* being the number fractions up to time *t*. Finally, assuming constant dose per fraction *d*, we can simplify the above equation:

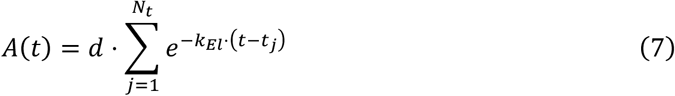

#### 3.3.2 Incidence of severe symptom: Markov chain model

We model PRO dynamics by employing an *n*-state minimal inhomogeneous continuous-time Markov chain (ICTMC) model with state 0 denoting symptom absence and state *n* − 1 denoting severe symptom (Figure 2C). The symptom intensity probability distribution *P*(*s* ∣ *t*) for landmark time *t* and time horizon *s* is defined as the solution to the Kolmogorov forward equation 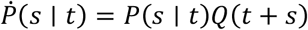 with transition rate matrix *Q*(*t*) (Figure 2D). We consider *Q*(*t*) of the following form:

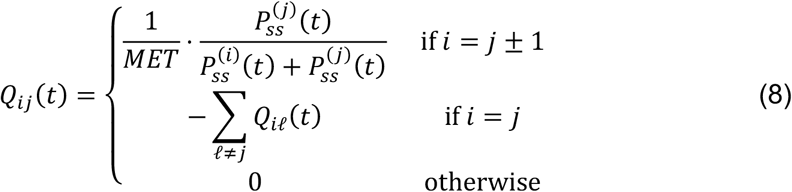

where *Q_ij_* (*t*) is the transition rate [day^-1^] from state *i* to state *j* at time *t*; *MET* [day] is the mean equilibrium time, which measures the strength of the Markovian property; and 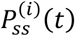is the steady state probability of being in state *i* at time *t* conditioned on parameters *θ*:

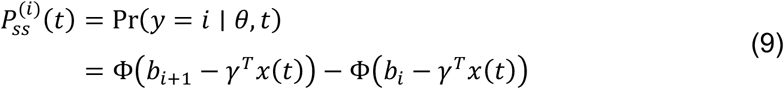

where Φ(*⋅*) is the standard normal cumulative density function; −∞ = *b*_0_≤ ⋯ ≤ *b_i_* ≤ ⋯ ≤ *b_n_* = ∞ are thresholds; and *γ* is a vector of effect sizes of 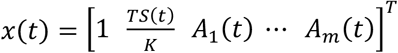 for *m* OARs. The form of *x*(*t*) is defined according to the OAR-symptom associations in Figure 1B.

The initial condition *P*(0 ∣ *t*) is set to *P_ss_* (*t*_0_) for the first observation at *t* = *t*_0_ and 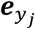 for subsequent observations *t_j_* > *t*_0_. Although not performed in this study, individual PROs can then be simulated by sampling *y_j_* from the distribution defined by *P*(*s*_*j*−1_ ∣ *t*_*j*−1_ and resetting the initial condition to the sampled value. We furthermore compute the mean symptom severity ^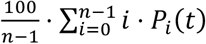^ *⋅ P_i_*(*t*), ranging from 0 (none) to 100 (severe) (Figure 2E). Finally, we compute the cumulative incidence of severe symptom (state *n* − 1) by considering it to be an absorbing state and setting *Q*_*n*−1_,_*n*−2_*t*) = 0 (Figure 2F).

### 3.4 Response and toxicity endpoints

To evaluate tumor response to treatment, we use 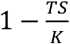 as a surrogate to model tumor control probability (TCP). To evaluate toxicity, we model symptom-specific normal tissue complication probability (NTCP) using the cumulative incidence of severe symptom. We then compute total NTCP as the cumulative incidence of experiencing any severe symptom: *NTCP* = 1 − ∏_*s*∈*symptoms*_ (1 – *NTCP_s_*). Finally, tumor response and toxicity are considered in tandem by computing the uncomplicated tumor control probability (UTCP): *UTCP* = *TCP ⋅* (1 − *NTCP*). We take our time endpoint the end of treatment on day 49.

### 3.5 Radiation plans: dose, OAR sparing, and fractionation

In this study, we define RT plans based on 3 characteristics: 1) total biologically effective dose (BED) delivered to the tumor; 2) OAR sparing; and 3) fractionation regimen. For simplicity, we take a coarse-grained approach and consider the equivalent uniform dose (EUD) delivered to the tumor and each OAR respectively, ignoring intra-organ heterogeneity (e.g., dose-volume histograms (DVH), 3D dose maps, functional subunits (FSU)).

OAR sparing is characterized by the tumor coverage (i.e., PTV isodose line; Figure 3A) and the dose delivered to each OAR. For simplicity, we assume that the dose delivered to each OAR is proportional to dose delivered to the tumor (Figure 3B). However, sparing OARs at the cost of reduced tumor coverage will sacrifice tumor response even for the same total EUD delivered to the tumor due to intra-tumoral heterogeneity. For example, RT plans with more OAR sparing and less tumor coverage are going to target the more hypoxic, radioresistant tumor cells in the necrotic core, whereas RT plans with less OAR sparing and greater tumor coverage are going to target the more vascularized, radiosensitive tumor cells in the proliferative edge.

**Figure 3.**
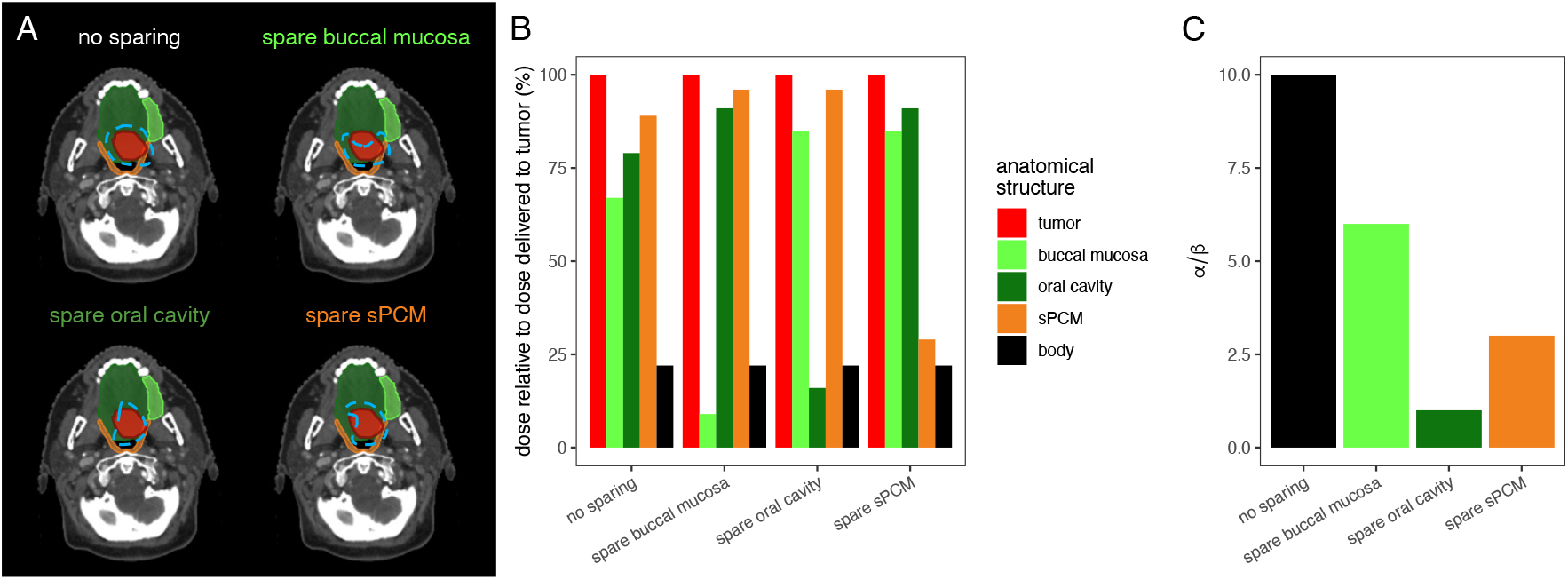
RT plans as characterized by OAR sparing. OAR sparing is characterized by: **A)** tumor coverage (i.e., PTV isodose line), **B)** proportions of dose delivered to each OAR relative to dose delivered to the tumor, and **C)** radiosensitivity via ^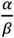^.

Therefore, we model the effect of OAR sparing on tumor response by lowering the linear radiosensitivity term *α*, which in turn lowers ^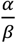^ (Figure 3C).

Three fractionation regimens are considered here: standard fractionation (1 fraction per day, Monday–Friday), hyper-fractionation (2 fractions per day, Monday– Friday), and hypo-fractionation (2 fraction per week (Monday, Wednesday)). In all cases, RT is administered over 7 weeks.

### 3.6 Model parameterization

Following Zahid, et al., we set tumor growth rate *k_g_* = 0.07 day^-1^, radiosensitivity parameters *α* = 0.09 Gy^-1^, *β* = 0.009 Gy^-2^, and ^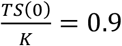^.^30^ Because we consider tumor size relative to carrying capacity ^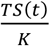^ as a covariate for the toxicity model, rendering *TS*(*t*) irrelevant, we set *K* = 1. Other model parameters were selected to qualitatively recapitulate the motivating data (Figure 1C–D) and are presented in Tables 1–3. We estimate the proportions of dose delivered to each OAR relative to dose delivered to the tumor for the maximum tumor control RT plan from Table 1 in Van den Bosch, et al. (Figure 3B)

**Table 1.** Tumor response parameters.

| parameter | description | unit | OAR sparing |  |  |  |
| --- | --- | --- | --- | --- | --- | --- |
|  |  |  | no sparing | spare buccal mucosa | spare oral cavity | spare sPCM |
| $TS_0$ | tumor size at treatment initiation | — | 0.9 | 0.9 | 0.9 | 0.9 |
| $k_g$ | tumor growth rate | day<sup>-1</sup> | 0.07 | 0.07 | 0.07 | 0.07 |
| $K$ | carrying capacity | — | 1 | 1 | 1 | 1 |
| $\alpha$ | linear radiosensitivity | Gy<sup>-1</sup> | 0.09 | 0.054 | 0.009 | 0.027 |
| $\beta$ | quadratic radiosensitivity | Gy<sup>-2</sup> | 0.009 | 0.009 | 0.009 | 0.009 |

**Table 2.** Absorbed dose decay rate by anatomical structure.

| parameter | description | unit | anatomical structure | value |
| --- | --- | --- | --- | --- |
| $k_{El}$ | radiation decay rate | day<sup>-1</sup> | buccal mucosa | 0.03 |
|  |  |  | oral cavity | 0.03 |
|  |  |  | PCM sup | 0.03 |
|  |  |  | body | 0.01 |

**Table 3.** Symptom parameters.

| parameter | description | unit | anatomical structure | symptom |  |  |  |
| --- | --- | --- | --- | --- | --- | --- | --- |
|  |  |  |  | oral pain | dysphagia | weight loss | tube feeding |
| $\gamma$ | effect size | — | intercept | -1.05 | -0.14 | -0.70 | 0.09 |
|  |  | — | tumor | 1 | 0 | 1 | 1 |
|  |  | Gy<sup>-1</sup> | buccal mucosa | 0.05 | 0 | 0 | 0.04 |
|  |  | Gy<sup>-1</sup> | oral cavity | 0.06 | 0.02 | 0 | 0.04 |
|  |  | Gy<sup>-1</sup> | PCM sup | 0 | 0.05 | 0.04 | 0.06 |
|  |  | Gy<sup>-1</sup> | body | 0 | 0 | 0.06 | 0.03 |
| $b$ | threshold | — | | [-0.18, 1.21, 2.19] | [-0.39, 0.73, 1.72] | 1.29 | 2.37 |
| $MET$ | mean equilibrium time | day | | 14 | 14 | 48 | 48 |

## 4 Results

### 4.1 Mathematical model recapitulates patient symptom dynamics

The developed mathematical model was able to recapitulate dynamics of the symptom intensity probability distribution observed in the published dataset (Figure 4A). Furthermore, the model was able to explain various features in the mean symptom intensity (Figure 4B): 1) a transient reduction of a subset of symptoms for 1–2 weeks, followed by 2) an acute exacerbation of symptoms throughout the rest of RT, followed by 3) a long-term relaxation of symptoms to below baseline levels.

**Figure 4.**
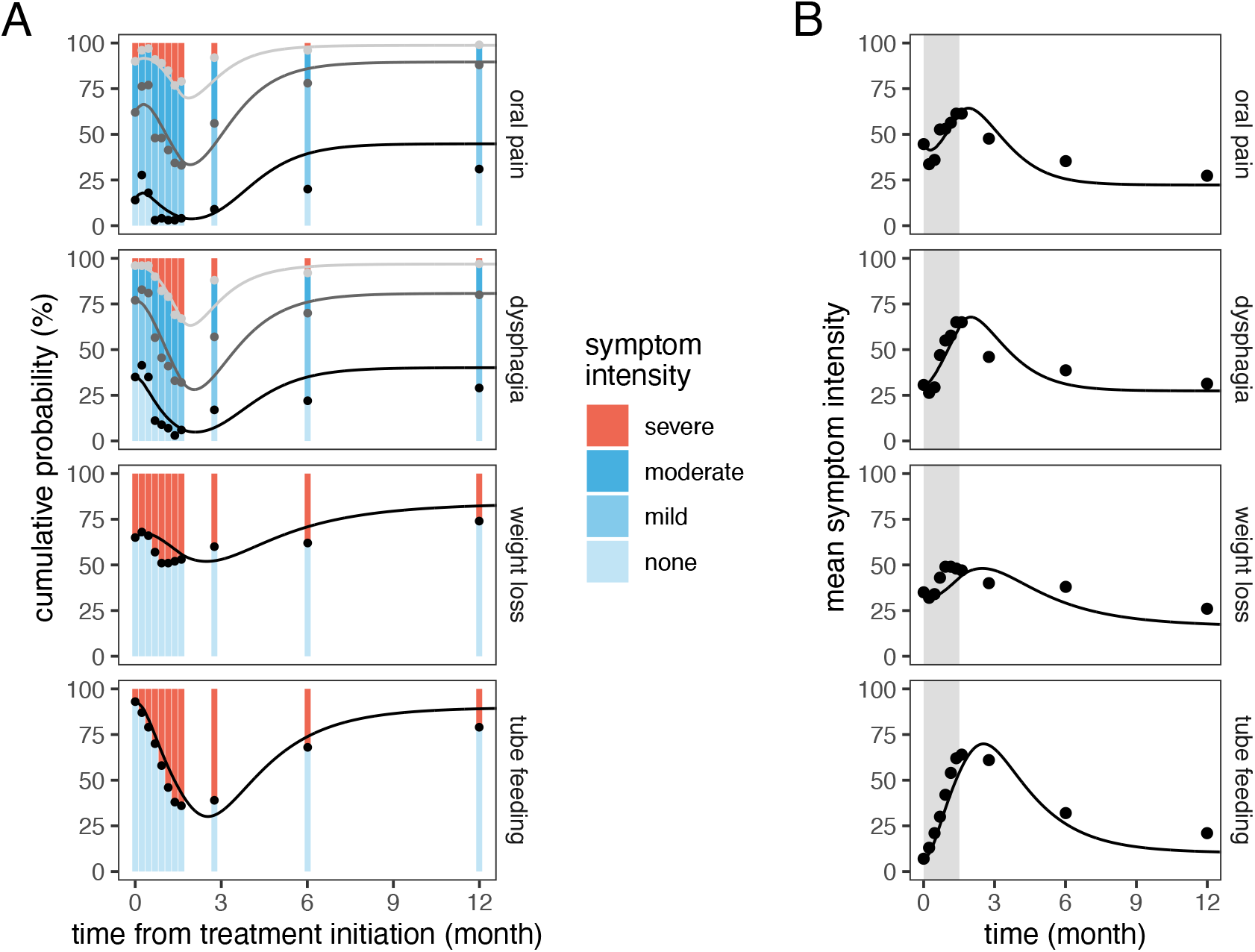
Mathematical model recapitulates published population-level PRO dynamics. **A)** Symptom intensity probability distribution. Black, dark gray, and light gray curves refer to probability of no, mild, and moderate symptom for oral pain and dysphagia. Black curve refers to no symptom for weight loss and tube feeding. **B)** Mean symptom intensity, ranging from 0 (none) to 100 (severe).

### 4.2 Tumor response and toxicity dynamics by dose, OAR sparing, and fractionation

As expected, tumors respond more with higher BED (Figure 5A). OAR sparing weakens the tumor response, due to lower radiosensitivity of the irradiated tumor.

**Figure 5.**
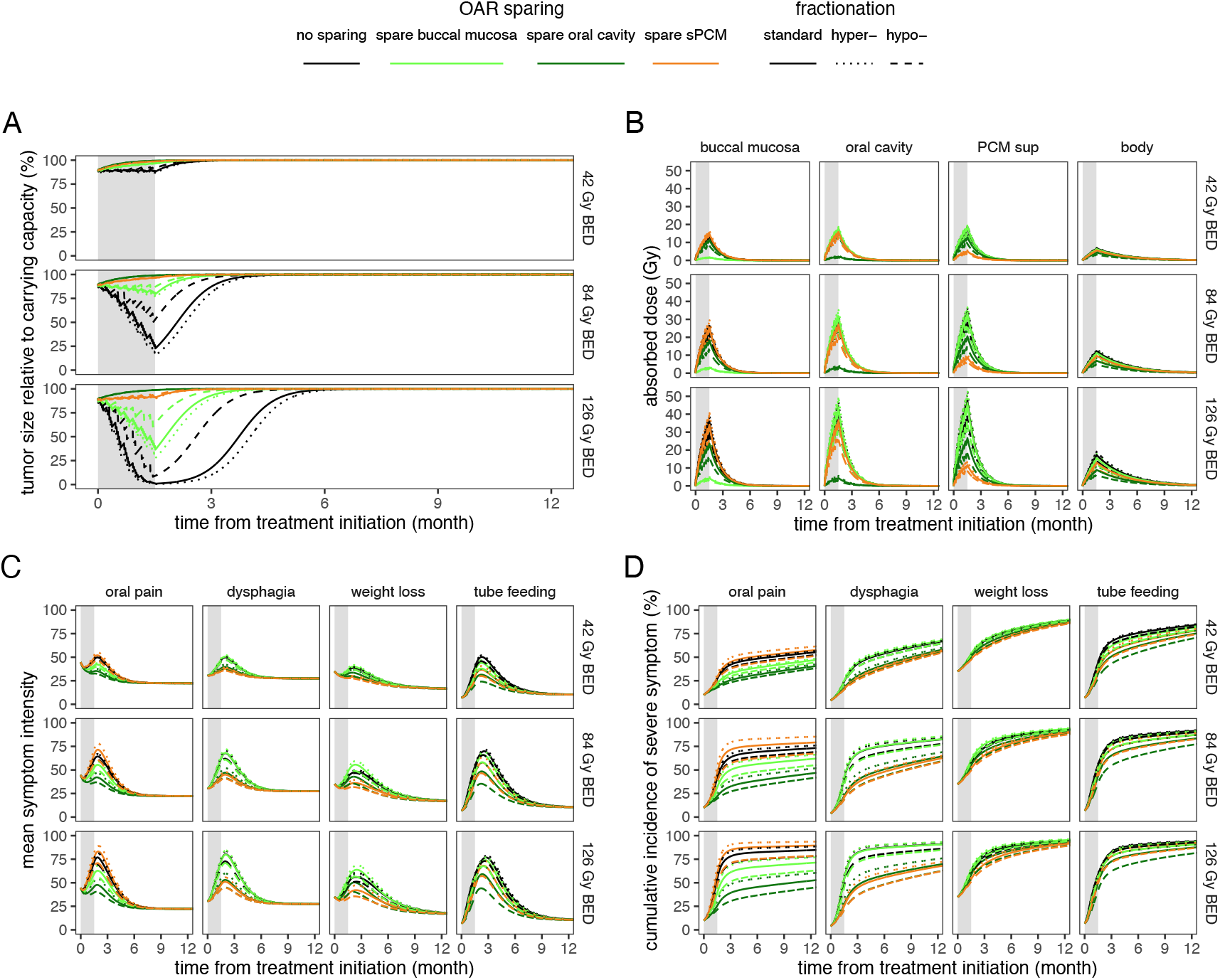
Tumor response and toxicity dynamics by total dose to tumor, OAR sparing, and fractionation. **A)** Tumor response dynamics. **B)** Absorbed dose kinetics for each OAR. **C)** Mean symptom intensity, ranging from 0 (none) to 100 (severe). **D)** Cumulative incidence probability of severe symptom. The gray region indicates 7 weeks of RT. For reference, 42, 84, 126 Gy BED are equivalent to 1.08, 2.00, 2.81 Gy per fraction for 35 fractions with an ^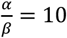^.

Hyper-(hypo-) fractionation strengthens (weakens) tumor response compared to standard fractionation. Notably, while high BED leads to near tumor eradication at the end of RT for all fractionation regimens, small differences in the tumor size become magnified in the post-RT tumor regrowth phase until saturation at carrying capacity.

Again, as expected, higher BED to the tumor leads to higher absorbed dose to OARs (Figure 5B). OAR sparing has variable effects on absorbed dose to OARs, which get magnified with higher BED. For example, sparing the buccal mucosa leads to less absorbed dose to the buccal mucosa but higher absorbed dose to the oral cavity and sPCM compared with the maximum tumor control plan. Hyper-(hypo-) fractionation leads to small increase (reduction) in absorbed dose compared with standard fractionation, which, like in the case of OAR sparing is magnified with higher absorbed dose and BED.

Again, as expected, higher BED to the tumor leads to higher symptom intensity and incidence of severe symptom (Figure 5C–D). As in the case of absorbed dose, OAR sparing has variable effects on symptom dynamics, which get magnified with higher BED. For example, sparing the buccal mucosa leads to less oral pain, but greater dysphagia and weight loss compared with the maximum tumor control plan. Hyper-(hypo-) fractionation leads to increase (reduction) in symptoms compared with standard fractionation, which is magnified with higher BED.

Finally, tumor response (TCP) and toxicity (NTCP) endpoints—modeled as tumor size relative to carrying capacity and incidence of severe symptom at the time endpoint, respectively—with respect to BED to the tumor, OAR sparing, and fractionation are shown in Figure 6A.

**Figure 6.**
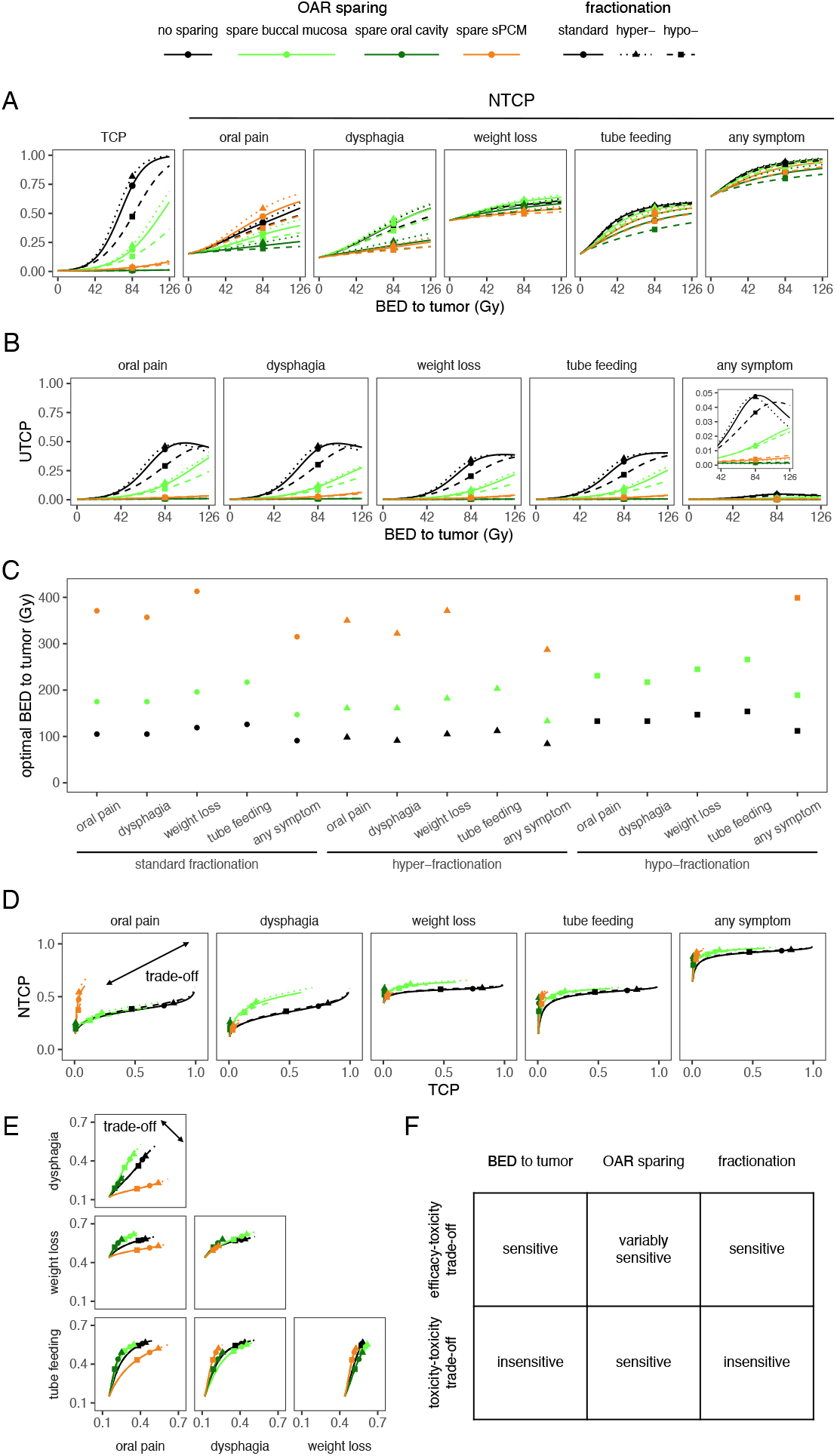
Tumor response and toxicity endpoints and their trade-offs by total dose to tumor, OAR sparing, and fractionation. **A)** Tumor control probability (TCP) and normal tissue complication probabilities (NTCP) by symptom. **B)** Uncomplicated tumor control probabilities (UTCP). **C)** Optimal total dose to tumor by UTCP. Sparing the oral cavity led to optimal BED much greater than 400 Gy (not shown). **D)** Response-toxicity trade-offs. **E)** Toxicity-toxicity trade-offs. **F)** Summary of effects of total dose to tumor, OAR sparing, and fractionation on response-toxicity and toxicity-toxicity trade-offs. Points denote 84 Gy BED to the tumor in all panels.

### 4.3 Dose finding to balance efficacy and toxicity

We consider tumor response and toxicity in tandem by computing the UTCP with respect to BED to the tumor, OAR sparing, and fractionation (Figure 6B). From here, we compute the optimal BED to the tumor, taken as the BED which maximizes UTCP (Figure 6C). For standard fractionation with maximum tumor control, the optimal BED was 91 Gy to achieve a UTCP of 4.8% for any symptom. This equates to a fractional dose 2.14 Gy, consistent with the standard of care. Hyper-(hypo-) fractionation decreases (increases) the optimal BED to 84 (112) Gy—or 1.08 (5.25) Gy per fraction— to achieve an UTCP of 4.7 (4.3) % for any symptom. OAR sparing drastically increases the optimal BED as the irradiated tumor is more radioresistant and toxicities are better tolerated. For example, for sparing the sPCM, the optimal BED is 315, 287, and 399 Gy to achieve an UTCP of 1.9, 1.4, and 2.8 % for any symptom for standard, hyper-, and hypo-fractionation, respectively. These BEDs equate to fractional doses of 3.91, 2.31, and 7.87 Gy, respectively.

### 4.4 Response-toxicity and toxicity-toxicity trade-offs

As expected, there is an observable trade-off between tumor response and off-target toxicities with respect to BED (Figure 6D). For example, a very low dose may result in negligible toxicity (low NTCP), but at the cost of marginal tumor response (low TCP). Alternatively, a very high dose may result in complete tumor regression (high TCP), but at the cost of high toxicity (high NTCP). Sparing a given OAR has the effect of lowering the tumor response (low TCP) while exhibiting a variable effect on toxicity. For example, compared to the maximum tumor control plan, sparing the sPCM lowers dysphagia but at the cost of reduced tumor response. However, sparing the sPCM does not exhibit this trade-off for oral pain. In fact, oral pain is increased while TCP is reduced. Fractionation also appears to induce a response-toxicity trade-off. Compared with standard fractionation, hyper-fractionation increases TCP at the cost of increasing NTCP, whereas hypo-fractionation reduces NTCP at the cost of reducing TCP.

Trade-offs between toxicities are sensitive solely to OAR sparing and insensitive to BED and fractionation (Figure 6E). For example, sparing the sPCM assuages severe dysphagia and weight loss. However, this comes at the cost of greater risk of severe oral pain. By contrast, increasing (decreasing) BED exacerbates (assuages) all symptoms. Likewise, hyper-(hypo-) fractionation exacerbates (assuages) all symptoms, compared with standard fractionation. Figure 6F summarizes the effects of BED to the tumor, OAR sparing, and fractionation on response-toxicity and toxicity-toxicity trade-offs.

## 5 Discussion

Here, we developed a mathematical model describing graded symptom dynamics and exploring response-toxicity and toxicity-toxicity trade-offs of RT in patients with HNCs. To do so, we modeled long-term normal tissue damage and repair using absorbed dose kinetics as a surrogate. However, it remains a question as to the interpretation of such a quantity. One possible biological interpretation of this concept involves the cascading role of localized ionizing radiation beams to create reactive oxygen species, which in turn leads to lethal DNA damage (e.g., double strand breaks)^31^ in concert with bystander effects.

We also took a coarse-grained approach in our model of dose across OARs by scaling proportionally to the dose delivered to the tumor. Future iterations of the developed model should leverage more fine-grained spatial, anatomical, and physiological information from CT scans. For example, some of this heterogeneity could be leveraged using 3D dose maps and dose-volume histograms (DVH).

An alternative approach may be to model normal tissue damage directly by employing an LQ formulation for normal healthy tissue. In this case, if we assume negligible impact of tumor on the surrounding normal tissue homeostasis, then we could set the initial condition at or near steady state. Following normal tissue complication by RT, we could model tissue repair mechanisms and return to homeostatic conditions.

Finally, we could employ a Markov chain model with normal tissue deviation from homeostasis, rather than absorbed dose kinetics, as a time-varying covariate.

In the case of many PRO items, another potential avenue for further model development is dimensionality reduction via a dynamic factor model, whereby latent constructs (e.g., cancer-related, treatment-related, and comorbidity-related symptoms) are inferred to explain time-to-event outcomes.^23^ By introducing latent constructs, we could directly model their dynamics using an Ornstein-Uhlenbeck process,^23^ rather than indirectly from the Markov chain model for observed item responses via the dynamic factor model.

Many of the patients in the motivating data were treated by RT with or without concurrent chemotherapy (e.g., cisplatin, carboplatin), which acts as a radiosensitizer.^33^ However, this benefit comes at the cost of toxicities, including high-grade dysphagia.^34^ Furthermore, RT has been found to modulate the pharmacokinetics of cisplatin.^35^ Future iterations of the model should account for these phenomena. However, not every patient is eligible for concurrent chemotherapy. Ineligible patients may instead receive EGFR inhibitors (e.g., cetuximab).^33^ Other alternative therapeutics involve symptom management (e.g., prescription mouth wash, opioids, dietician, speech pathologist, psychologist). Furthermore, dose (de-) escalation,^36^ adaptive re-planning,^37,38^ as well as alternating between RT plans if one symptom is predicted to become too intolerable at a certain point in time to spare normal tissue while trying to maintain as much tumor response as possible. Future iterations of the model should allow for comparison of the above alternative therapeutic strategies.

The simulations performed in this study support our conceptual framework of leveraging mathematical modeling to find doses that balance response-toxicity and toxicity-toxicity trade-offs in head and neck cancer treated with RT. We envision that future iterations of the model developed here will aid clinicians in dose-finding and selecting a RT plan that will optimize trade-offs between target efficacy and toxicity endpoints selected by the patient in consultation with their healthcare providers.^22^

While the selected absorbed dose and toxicity model parameters were able to recapitulate published population-level PRO data, they were chosen by experimentation and visual inspection. Future implementations of this model will require careful calibration to patient data using either frequentist (e.g., maximum likelihood estimation (MLE)) or Bayesian (e.g., sampling by Hamiltonian Monte Carlo (HMC) simulations^32^) approaches. To do this, we will need to account for inter-patient dose heterogeneity to fit population-level longitudinal data.

Overall, the mathematical model presented in this study addresses some of the limitations of current NTCP models by explaining graded toxicity dynamics. By integrating both tumor control and quality of life considerations into a singular model, recommendations of dose, OAR sparing, and fractionation can be made. Future iterations of the model could aid clinicians in RT dose-finding and selecting a RT plan that will optimize tumor control and patients’ quality of life.

## Notes

Conflict of Interest: The authors have no relevant conflicts of interest to disclose.

### Competing Interest Statement

The authors have declared no competing interest.

### Summary of Updates

RT plans have been expanded to include both OAR sparing and fractionation. "Radiation exposure" has been recast as "absorbed dose kinetics." We shifted emphasis more onto graded NTCP dynamics. Finally, the author list has been updated.

